# A modular high-density 294-channel μECoG system on macaque vlPFC for auditory cognitive decoding

**DOI:** 10.1101/768127

**Authors:** Chia-Han Chiang, Jaejin Lee, Charles Wang, Ashley J. Williams, Timothy H. Lucas, Yale E. Cohen, Jonathan Viventi

## Abstract

**OBJECTIVE:** A fundamental goal of the auditory system is to parse the auditory environment into distinct perceptual representations. Auditory perception is mediated by the ventral auditory pathway, which includes the ventrolateral prefrontal cortex (vlPFC) late. Because large-scale recordings of auditory signals are quite rare, the spatiotemporal resolution of the neuronal code that underlies vlPFC’s contribution to auditory perception has not been fully elucidated. Therefore, we developed a modular, chronic, high-resolution, multi-electrode array system with long-term viability.

**APPROACH:** We molded three separate μECoG arrays into one and implanted this system in a non-human primate. A custom 3D-printed titanium chamber was mounted on left hemisphere. The molded 294-contact μECoG array was implanted subdurally over vlPFC. μECoG activity was recorded while the monkey participated in a “hearing-in-noise” task in which they reported hearing a “target” vocalization from a background “chorus” of vocalizations. We titrated task difficulty by varying the sound level of the target vocalization, relative to the chorus (target-to-chorus ratio, TCr).

**MAIN RESULTS:** We decoded the TCr and the monkey’s behavioral choices from the μECoG signal. We analyzed decoding capacity as a function of neuronal frequency band, spatial resolution, and time from implantation. Over a one-year period, we were successfully able to record μECoG signals. Although we found significant decoding with as few as two electrodes, we found near-perfect decoding with ∼16 electrodes. Decoding further improved when we included more electrodes. Finally, because the decoding capacity of individual electrodes varied on a day-by-day basis, high-density electrode arrays ensure robust decoding in the long term.

**SIGNIFICANCE:** Our results demonstrate the utility and robustness of high-resolution chronic µECoG recording. We developed a new high-resolution surface electrode array that can be scaled to cover larger cortical areas without increasing the chamber footprint.

## 1. Introduction

Real-world hearing is a complex computational problem for two main reasons: (1) auditory stimuli can simultaneously change along many dimensions, such as loudness and location, and (2) stimuli of interest (e.g., the voice of a friend) are often mixed together with other environmental sounds (e.g., the sound of frothing milk) [1, 2]. Thus, for a listener to interact efficiently with their environment, they must be able to selectively attend to stimuli of interest and importance (e.g., your friend’s voice), while simultaneously ignoring extraneous stimuli (i.e., the frothing milk) [2-7].

A classic test of real-world hearing is the detection of an auditory stimulus that is embedded in a noisy background [8-12]. For example, normal-hearing listeners can readily detect and hear a friend’s voice at a party, even when the background is noisy due to the voices of other speakers, music, and clinking glasses. This is often referred to as the “cocktail-party problem” or “hearing in noise” and is one of the fundamental challenges of the auditory system [13].

Although the neuronal correlates of hearing in noise have been studied acutely in a variety of model species and using many different techniques and stimulus paradigms [8, 9, 14-24], these prior works focused on early cortical centers and neglected the contribution of later regions, like the ventrolateral prefrontal cortex (vlPFC). The vlPFC is part of the of the ventral auditory pathway, which is widely considered to play a substantial role in auditory perception and cognition [25].

In addition to not understanding vlPFC’s contribution to fundamental tasks like hearing-in-noise, we only have a rudimentary understanding of how information is encoded in populations of vlPFC neurons [21, 24, 26-42]. Because of this poor understanding, we do not have a good understanding of the spatiotemporal resolution of the neuronal code that underlies vlPFC’s contribution to perception. The topographic and temporal scale of these neuronal populations has not been tested because, until recently, it was not possible to record from large numbers of brain sites simultaneously [43]. Even today, such large-scale recordings in the cortical auditory system are still very rare.

To overcome some of the challenges of recording from neuronal populations and to do so chronically [44-46], we demonstrate the feasibility of a new micro-electrocorticographic (μECoG) array that provides, to our knowledge, the highest electrode density and channel count in any non-human primate study (Table 1). The advantage of this technique is that the array is assembled out of smaller sub-arrays that are combined into one larger one. Because of this technique, it is possible to tailor the size, shape, and electrode density for the particular brain target. This design also permits the recording chamber to house electronics that scale vertically as channel count increases, giving more room for electrode-interface connections. A water-tight recording chamber design enables electronics to be permanently implanted on the animal, to allow for future channel-count scaling. We show that μECoG signals from this array contain multiple sources of information that relate to a monkey’s performance on a hearing-in-noise task. We further show that we had almost near-perfect decoding accuracy after 1-year of implantation.

**Table 1.** µECoG comparison table. Comparison between this work and other µECoG studies.

| Studies | Year | Subject | Number of Electrodes | Spacing (mm) | Coverage (mm<sup>2</sup>) | Density (Sites/mm<sup>2</sup>) |
| --- | --- | --- | --- | --- | --- | --- |
| Hollenberg et al. [84] | 2006 | Rat | 64 | 0.75 | 36 | 1.78 |
| Benison et al. [85] | 2007 | Rat | 256 | 0.5 | 64 | 4.00 |
| Molina-Luna et al. [86] | 2007 | Rat | 72 | 0.64 | 27 | 2.67 |
| Kim et al.[87] | 2007 | NHP | 56 | 1 | 36 | 1.56 |
| Rubehn et al. [88] | 2009 | NHP | 252 | 2 | 2100 | 0.12 |
| Ledochowitsch et al. [89] | 2011 | Rat | 256 | 0.5 | 64 | 4.00 |
| Besle et al. [90] | 2011 | Human | 127 | 10 | 12700 | 0.01 |
| Viventi et al. [91] | 2011 | Cat | 360 | 0.5 | 90 | 3.60 |
| Pasley et al. [92] | 2012 | Human | 64 | 4 | 1000 | 0.06 |
| Khodagholy et al. [93] | 2015 | Rat | 64 | 0.1 | 1 | 64.00 |
| Escabi et al. [94] | 2014 | Rat | 196 | 0.25 | 12.25 | 16.00 |
| Hotson et al. [95] | 2016 | Human | 128 | 3 | 1152 | 0.11 |
| Kellis et al. [96] | 2016 | Human | 16 | 1 | 16 | 1.00 |
| Khodagholy et al. [97] | 2016 | Human | 240 | 0.23 | 840 | 0.29 |
| Kaiju et al. [98] | 2017 | NHP | 96 | 0.7 | 4704 | 0.02 |
| **This work** | **2019** | **NHP** | **294** | **0.61** | **114.4** | **2.57** |

## 2. Materials and Methods

### 2.1. Electrode

Our modular 294-channel µECoG electrode array was built from three separate 98-channel arrays. These arrays were fabricated using microfabrication methods in a cleanroom environment at the Shared Materials Instrumentation Facility at Duke University. The full array was composed of 294 electrode contacts (229-µm diameter) with an inter-electrode pitch of 610 µm (center-to-center). A local reference electrode (38.1-mm long × 200-µm wide) surrounded the contacts (Figure 1a). This yielded a total sensing area of 10.4 × 11 mm^2^ with a density ∼2.6 sites/mm^2^.

**Figure 1.**
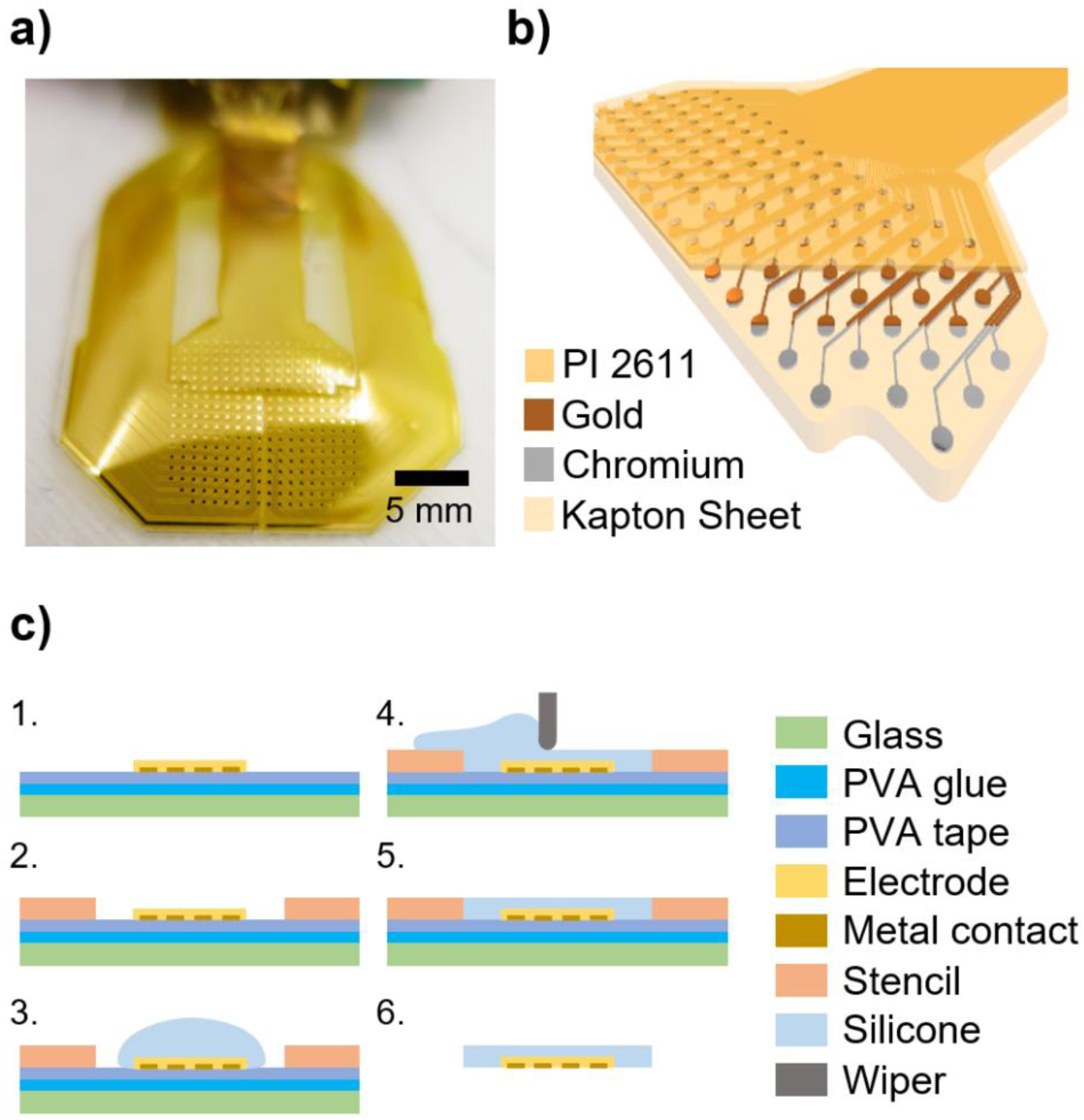
Electrode assembly and fabrication. **(a)** Molded electrode array with a high-resolution sensing area (10.4 × 11 mm^2^) including 294 electrode channels. **(b)** A 3D cross sectional schematic of the electrode array. The electrodes were fabricated on top of a 25-µm Kapton Polyimide sheet. 20 nm of chromium and 250 nm of gold were E-Beam deposited to form the electrode contacts and traces. 6 µm of polyimide (PI 2611) was spin coated onto the device to serve as the top encapsulation layer. The electrode contacts and connector openings were etched open using RIE. **(c)** Illustrations of the electrode molding process. 1. PVA glue was applied to a glass slide to secure PVA tape. Electrode arrays were then laminated onto the PVA tape in order to temporarily hold the electrode sub-arrays for alignment. The PVA tape also prevented silicone from accidently covering up the electrode contacts during the molding process. 2. A stencil was placed onto the assembly to check the electrode alignment and to set the thickness of the silicone molding. 3. Well-mixed and de-gassed silicone was poured onto the mold. 4. A wiper removed the excessive silicone. 5. The silicone cured at room temperature for 24 hours. 6. The assembly was rinsed under tap water to remove the PVA tape and release the molded array.

#### 2.1.1. Fabrication

The electrode stack-up is shown in Figure 1b. Similar to our previous work [47], the base electrode substrate consisted of a 25-µm Kapton® Polyimide sheet (Fralock, Inc., Valencia, CA), which was manually laminated onto a glass slide coated with cured polydimethylsiloxane (PDMS) (Sylgard 184, Dow Corning, Midland, MI). Using an e-beam metal evaporator (CHA Industries E-Beam), 20 nm of Chrome (Cr) and 250 nm of Gold (Au) were deposited onto the Kapton sheet. S1813 positive photoresist (Shipley Microposit) was used as a positive mask to wet etch the Au and Cr layers (Gold Etch TFA, Cr Etchant 9057; Transene, Danvers, MA). Next, 6-µm of polyimide (PI 2611, HD Microsystems, Parlin, NJ) was spun onto the surface of the array and cured. The top polyimide layer was selectively etched using a Trion Phantom II reactive ion etcher (RIE). The electrodes were then removed from the glass substrate. Finally, a stiffener, which was composed of a 50-µm thick layer of pressure sensitive adhesive (3M467-PSA) and 153 µm of polyimide, was applied to the back of each 51-pin zero-insertion-force (ZIF) connector, two for each 98-channel array. Individual electrodes were screened by measuring their impedance (@ 1 kHz) in saline solution using a NanoZ system (White Matter LLC, Mercer Island, WA).

#### 2.1.2. Assembly (molding)

We used biocompatible medical-grade silicone molding (MDX4-4210, Dow Corning, Midland, MI), which is similar to silicone that we used previously as artificial dura [48, 49]. Using this silicone, we combined three separate electrode sub-arrays together to construct a single uniform high-density electrode array. This design resulted in a narrow cable entry point to the recording chamber (Figure 1a). Because the silicone also covered the sharp electrode edge of the thin polyimide, it reduced the possibility of damage to the brain during array placement.

The molding process is shown in Figure 1c. Electrodes were manually aligned on a glass slide with water-soluble (Polyvinyl alcohol, PVA) tape (3M 5414). The tape prevented the electrode contacts from being covered by silicone and maintained alignment during molding. We designed and fabricated custom molding and alignment stencils to ensure that the silicone-molded 294-channel µECoG array had a uniform shape and thickness. Once the silicone cured, the molded electrode array was released by rinsing the array under water. With only a 100-µm thick coating of silicone molding, the molded array remained flexible. The functionality of the array was bend tested to a radius of 2.5 mm using a glass rod (Supplement Fig. 1).

### 2.2. Chamber

Commercially available recording chambers typically require manual adjustments during surgery, such as bending the mounting legs, to secure the implant on the skull [50, 51]. These manual adjustments may take a lot of time during surgery, increasing the length of the procedure. These adjustments also create gaps between the mounting pedestal and skull. Because these recording chambers have a universal design, they may not fit tightly to the skull and consequently, may not fully integrate with the bone. If a chamber fails to integrate with the skull bone, it can come loose or break off, ending the experiment and endangering the animal [52].

Recent studies have utilized 3D printing to create custom chambers to address this challenge [53, 54]. However, these chambers were designed for penetrating microdrives rather than surface electrodes and were not designed to be watertight, which is essential for implants that house electronics and PCBs.

To address these challenges, we used a 3D-printed chamber to create a custom base curvature that sat seamlessly on the skull. The skull model was generated from a structural MRI study that was conducted prior to the monkey’s surgery [52]. The chamber structure was optimized to improve impact resistance, based on computer simulations. The chamber system was composed of three major components: 1) a 3D-printed custom titanium chamber to ensure a seamless fit between the skull and the implant, 2) a molded silicone rubber chamber wall (Sugru, FormFormForm Ltd., London, United Kingdom), and 3) a molded silicone gasket. The wall and the gasket worked together to form a watertight chamber.

#### 2.2.1. Identification of vlPFC

vlPFC was initially identified through structural MRI scans of the monkey’s brain [55-58]. vlPFC is dorsal of the inferior ramus of the arcuate sulcus. vlPFC neurons are functionally identified by their task-related auditory responses [41, 59, 60]. The left hemisphere was targeted to facilitate comparisons with previous findings [7, 41, 61-67].

#### 2.2.2. Skull Mapping

To create a custom base that fit the contour of the skull, we first extracted the portion of the skull MRI data that was located under and near the potential chamber location. We then performed a re-mesh step (MeshLab, Visual Computing Lab ISTI-CNR) that reduced the number of triangles in the 3D skull model from ∼1 million to ∼3000. This step balanced the costs associated with long processing times with the detail needed to produce a realistic model of skull curvature. Finally, we exported this simplified skull surface model to Autodesk Inventor (Autodesk Inc., San Rafael, CA) to complete the chamber design.

#### 2.2.3. Design and fabrication

With the simplified skull model, we were able to quickly iterate designs to optimize the structure and mounting strategy (Figure 2). We 3D printed low-cost plastic test samples, which allowed our surgeon to optimize the locations of the bone screws in advance. These low-cost test samples were made out of Polylactic Acid (PLA) via the common fused filament fabrication (FFF) method. The test samples also allowed test fitting and fine tweaks of the design before we submitted the design for titanium printing. We used finite element analysis software simulation tools (Autodesk Nastran, Autodesk Inc., San Rafael, CA) to optimize our design parameters (e.g., shape and wall thickness), which improved the chamber’s impact resistance.

**Figure 2.**
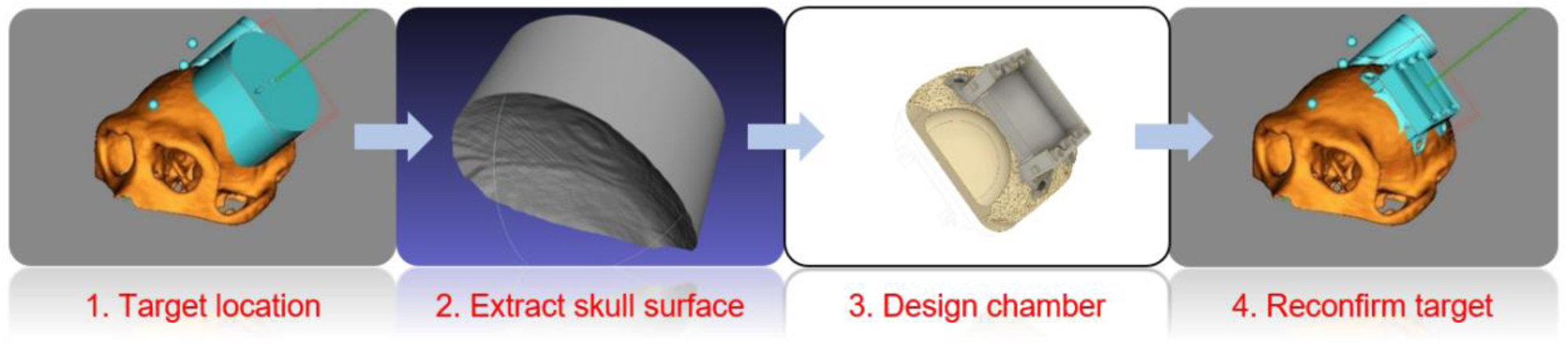
General procedure for creating the custom chamber base. 1. Structural MRI data determined the target location and the corresponding skull location in which to mount the chamber. 2. The target skull surface was extracted and simplified to reduce the curvature details and processing time. 3. The chamber was then iterated to optimize the design and mounting strategy. 4. The final chamber design was re-imported back into the targeting software environment to validate placement.

To simulate a worst-case scenario, instead of calculating the average impact force, we calculated the dynamic energy of a free-falling monkey at the moment when the chamber hits the ground From the work-energy principle, the maximum impact force is calculated as: 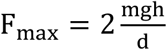, where m is the mass of the falling object in kilograms, g is the gravitational acceleration constant (9.8 m/s^2^), is the falling distance in meters, and is the impact distance in meters.

The impact force on a monkey’s chamber can be difficult to determine because it depends on how the chamber hits the ground, (e.g., which part of it hits the ground, the angle of impact, and if it was somehow protected). We chose an impact distance of 6 cm, along with a 1.2-m falling distance, and a monkey weight of 11.0 kg. This yields an impact force of ∼4300 N. Our simulation found that the maximum yield strength was ∼700 MPa (Figure 3), when the impact force occurred on the top of the chamber, which is below the yield strength limit of ∼900 MPa for titanium. Additional simulation results for impacts on the front, side, and corner of the chamber are shown in Supplement Fig 2.

**Figure 3.**
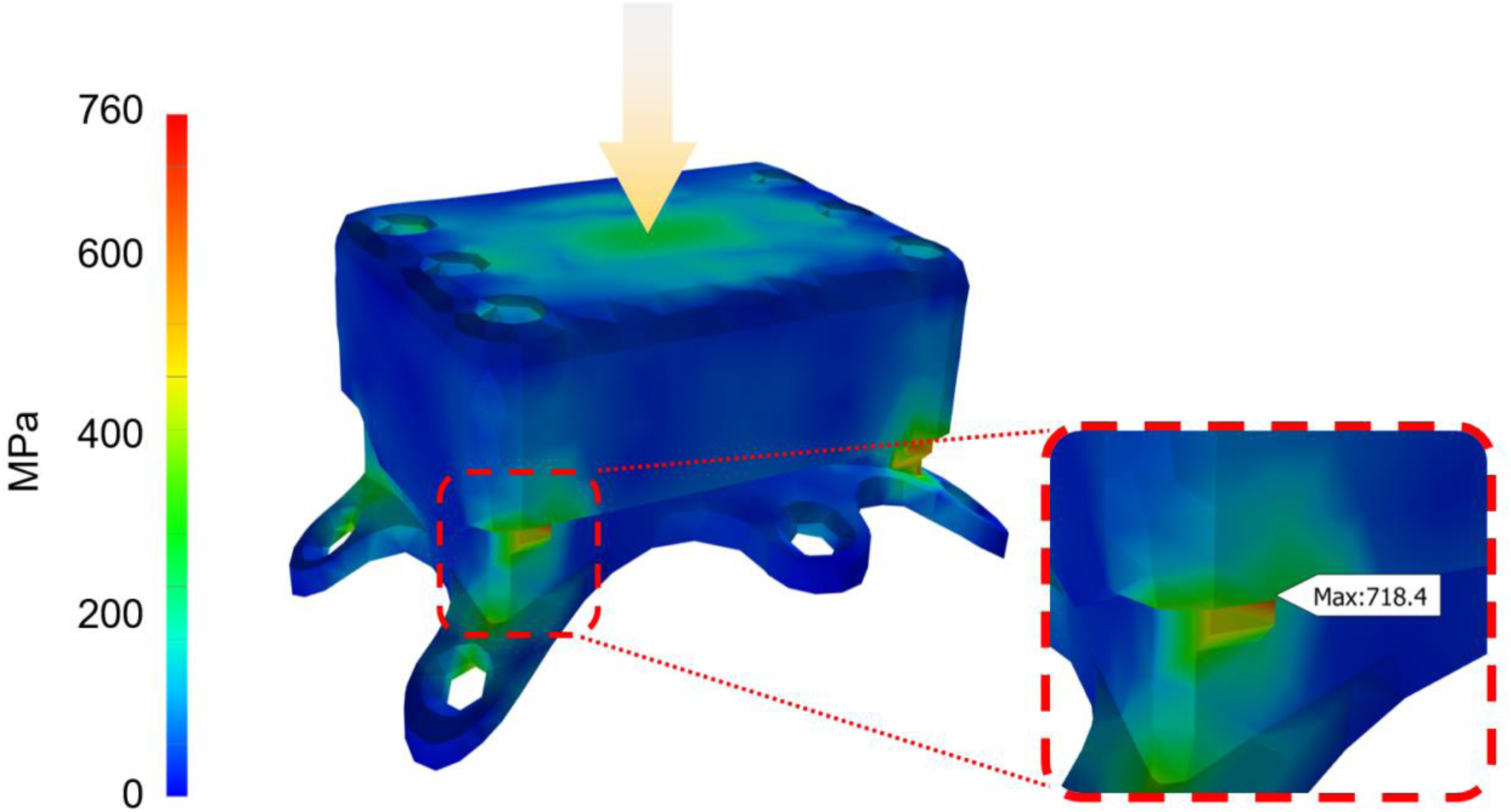
Mechanical simulation of free-fall impact force. We optimized the chamber structure to withstand the impact force of a free-falling NHP. The most vulnerable point of the chamber was the safety stopper tab (red dash square) which prevented the cap from being over-tightened. When an impact force was applied from the top surface (yellow arrow, ∼4300N), a maximum pressure of 718.4 MPa (maximum point identified by the simulation software) occurred in the tabs, which is below the yield strength limit of ∼900 MPa for titanium. The simulation results from impact forces in different directions are shown in Supplement Fig. 2. The impact force distribution in the chamber body over time is shown in movie M1.

Water resistance was another important design factor because we permanently attached the adapter PCBs to the chamber. To make the chamber water tight, we constructed: 1) a custom molded silicone rubber wall with a reinforcement stiffener and 2) a silicone molded gasket between the titanium chamber base and cap. The chamber stack-up is shown in Figure 4b.

**Figure 4.**
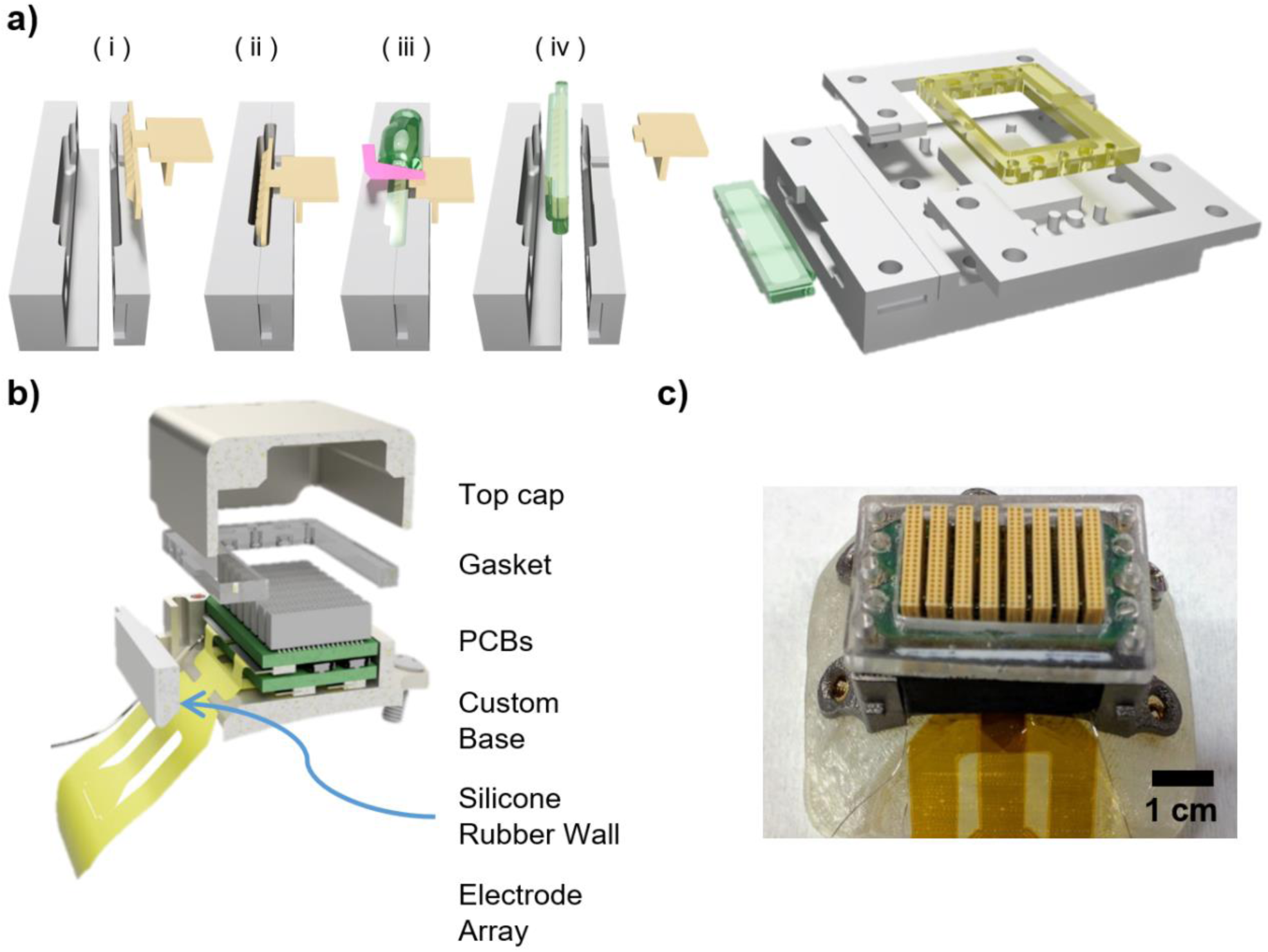
Chamber system assembly steps. **(a) Left.** Molding steps for the silicone rubber wall. (i) Demolding agent was applied to the aluminum mold. (ii) The mold was assembled and the stiffener piece was placed at the center of the mold. (iii) The mold was filled with silicone rubber (Sugru) and excess material was removed using a squeegee. (iv) After curing the mold in an oven (60°C, 4+ hours), the molded wall was removed from the aluminum mold and the alignment portion of the stiffener was broken off. **Right**. A 3D rendering of the aluminum mold that was used to mold the silicone rubber wall and the silicone gasket. The molding steps of the silicone gasket were similar to the silicone rubber wall but without the stiffener. Silicone (MDX4-4210) was used for the gasket. **(b)** Exploded 3D view of the chamber system. **(c)** The final chamber on top of a model skull. Detailed assembly steps are shown in movie M2.

The silicone rubber wall compressed and deformed around the electrode to form a watertight seal. A 3D-printed stiffener (DuraForm PA, 3D System) reinforced the silicone rubber wall and helped it to retain its shape under pressure. The reinforcement stiffener was printed using selective laser sintering. This printing method increased the surface roughness of the stiffener, which, in turn, improved the adhesion of the stiffener to the silicon rubber.

To seal the gap between the chamber base and cap, we designed a custom silicone (DuPont MDX4-4210) gasket. The gasket was compressed by the chamber cap to seal the chamber. The gasket was replaced if there was any sign of wear or damage.

To make the custom gasket, we first applied a thin layer of a medical-grade biocompatible mold-release agent (Duraglide MCC-DGF14A, MicroCare Corp., New Britain, CT) to the surface of the stencil mold. This facilitated removal of the molded gasket after it cured. We next placed well-mixed silicone in a desiccator for 30 min to remove the air bubbles trapped during mixing and then poured into the mold; we heated the mold to accelerate the curing process (60°C for 6+ hours). The silicone rubber wall was constructed in an analogous manner, using the same aluminum stencil mold. Detailed molding steps and the corresponding stencil molds for both parts are shown in Figure 4a.

We tested the water resistance of the chamber by placing a water-contact indicator strip (3M 5559) within a fully assembled chamber and soaking it in phosphate buffered saline (PBS; pH = 7.4; Quality Biological, MD) at room temperature. For visual confirmation, we also 3D printed (Transparent Resin, 3D Hubs, Amsterdam, Netherlands) a water clear chamber using the Polyjet method (Supplement Fig. 3). We found that, for both the titanium and clear chambers the indicator strip remained dry after several days of soaking.

After validation, the final design (resolution: ±100 µm) was 3D-printed in titanium (TiAl_6_V_4_) via direct metal laser sintering (3D Hubs). A picture of the final chamber is shown in Figure 4c.

### 2.3. Data acquisition system and *in vivo* recording

Typically, chamber size increases as the number of electrodes implanted increases. In this work, although having hundreds of electrodes, we minimized the chamber footprint by exploiting a modular stacking design that used off-the-shelf components. The final chamber size was reduced to 3.5 × 2.7 × 1.8 cm^3^, including mounting wings and weighed only 76 g including the PCBs, cap, gasket, screws, and all other hardware.

Within the chamber, two custom PCB interface boards were developed to connect the electrode array to the data-acquisition system. Four 51-pin ZIF (504070 series, Molex, IL) connectors connected the level-1 interface board with two of the 98-channel arrays. Two additional ZIF connectors connected the level-2 interface board with the third 98-channel array. The level-1 board was connected to the level-2 board through two 100-pin high-density stacking connectors (55909 series, Molex, IL), which passed 196 electrode signals along with multiple reference signals from the level-1 board to the level-2 board. The top of the level-2 board included eight Omnetics connectors (NSD-36-VV-GS) that interfaced with four 64-channel amplifier headstages (RHD 2164, Intan Technologies).

With a maximum of 256 channels recorded from the four 64-channel amplifiers, we shorted the remaining outer contacts (Supplement Fig. 4) on the adapter boards to create another local reference ring that could be used in data analysis. Future designs could incorporate the amplifier integrated circuits into each level PCB, enabling all 294 recording electrodes.

Digital data from the Intan amplifiers were transferred via thin and flexible SPI tethers, and logged with the OpenEphys system [68] at 20 kS/s. The programmable hardware filter settings in the Intan headstages were set to 0.1 Hz – 7.5 kHz. The system diagram is shown in Figure 5.

**Figure 5.**
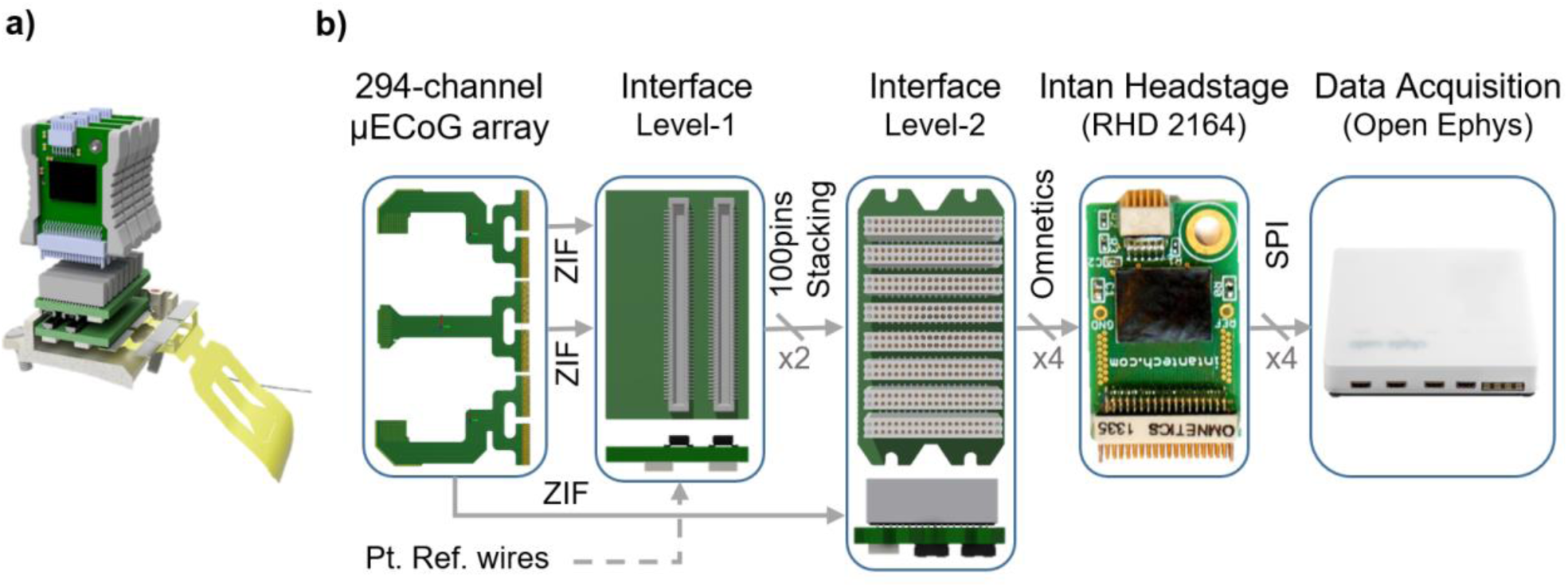
µECoG recording system. **(a)** A 3D rendering of the full system during recording. **(b)** System block diagram. Two of the electrode sub-arrays were connected to interface board level-1, and the third sub-array was connected to the interface board level-2. The two interface boards were connected via two 100-pin stacking connectors that passed 196 electrode channels and the local references. Eight Omnetics connectors were used to connect to the Intan headstages during recording. Side views of interface board levels-1 & -2 are also shown at the bottom of the panel. An OpenEphys controller recorded from the four Intan headstages.

### 2.4. Chamber implantation

Prior to implantation, the electrodes were connected to the interface PCBs, and the level-1 and level-2 PCBs were connected together. After this assembly step, the PCBs were conformally coated (Bondic, Laser Bonding Technology Inc., Ontario, Canada) to protect the circuits from potential water ingress and improve reliability. The electrode and PCB assemblies were gas sterilized (ethylene oxide; Duke University Medical Center).

The University of Pennsylvania Institutional Animal Care and Use Committee approved all of the surgical and experimental protocols. All surgical procedures were conducted using aseptic surgical techniques, during which the monkey was under general anesthesia.

In brief and following from our previous work [52], the scalp and the periosteum were incised along the midline. After stereotactically identifying the vlPFC [57], we used a piezoelectric drill (Synthes) with a small round cutting burr to perform a craniectomy. Next, after the dura was incised and reflected, the µECoG array (which was attached to the titanium chamber) was placed on the brain surface (Figure 6). The dura was then overlain and the calvarium was replaced. The recording chamber was attached to the skull via Ti bone screws; the screws were inserted through holes in the legs of chamber. The legs and edges of the craniectomy were protected with Geristore (DenMat). Finally, the cover was placed and secured on the recording chamber, and the skin edges were relaxed and sutured over the chamber.

**Figure 6.**
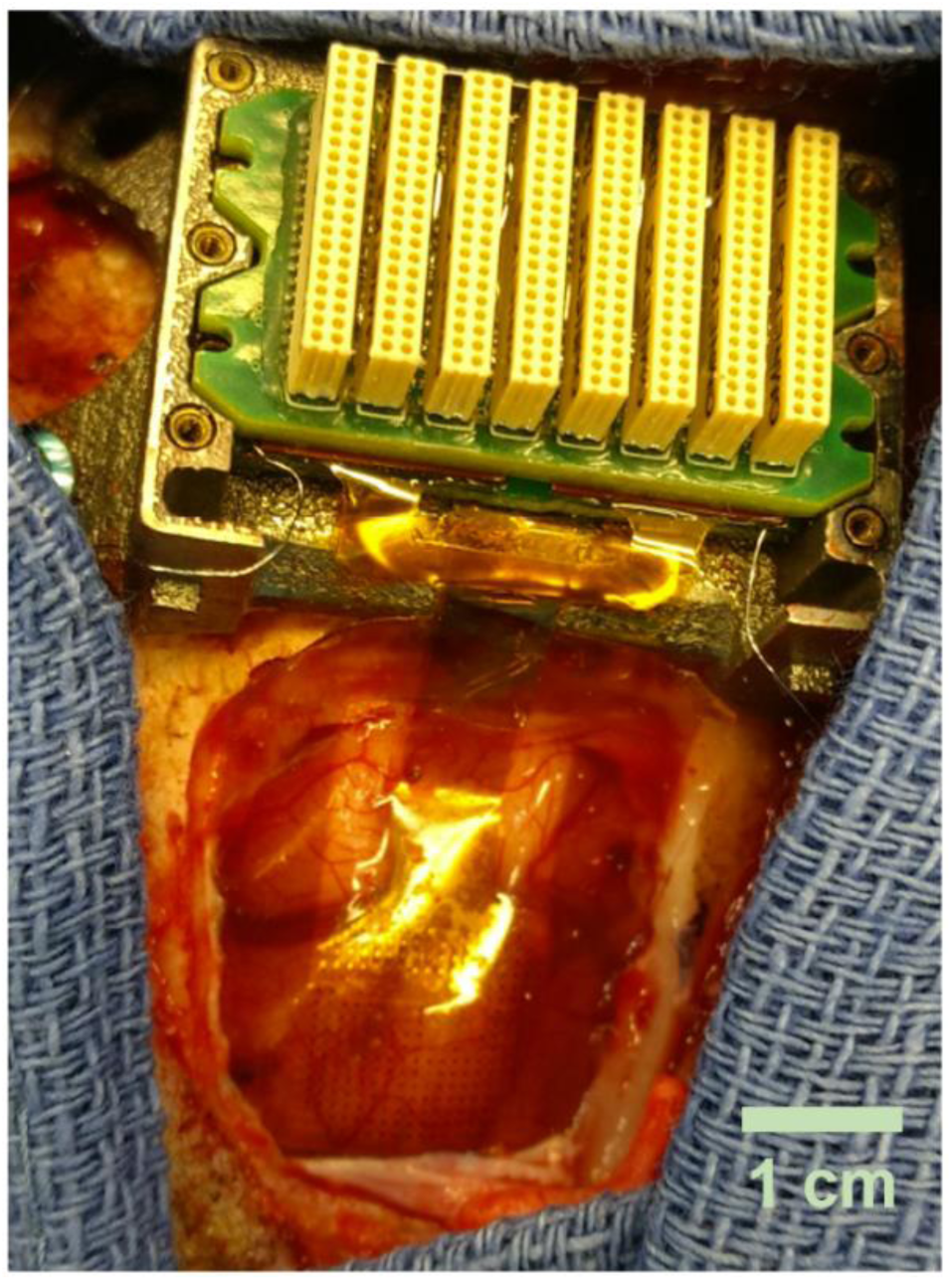
Image of the implanted µECoG array. At the bottom of the image, the craniectomy and the electrode array that is lying on the surface of the vlPFC is visible. This image was obtained prior to overlaying the dura on the array and brain surface. The chamber with the electronics is shown at the top of the picture.

### 2.5. Behavioral Task

The hearing-in-noise task tests a monkey’s ability to detect a <u>target vocalization</u> that is embedded in a <u>background chorus</u> of vocalizations. 400-700 ms after the monkey grasped a touch-sensitive lever, we presented a target vocalization that was embedded in a background chorus. Following onset of the target vocalization, the monkey had 600 ms to move the lever to report hearing the vocalization.

The target vocalization was a single exemplar of a *coo* (duration: 400 ms) that was recorded from an unknown conspecific [69, 70]. The background chorus was created by superimposing 30-40 different vocalizations, which were also from unknown conspecifics, at random times to create a 6600-ms stimulus; on a day-by-day basis, we created new tokens of this background chorus. We minimized the amplitude troughs of this stimulus mixture to reduce the possibility that the monkey could detect the target vocalization if it occurred within an amplitude trough of the background chorus [21, 71]. The sound level of the target vocalization was varied relative to the level of the background chorus (65 dB SPL): the <u>target-to-chorus ratio</u> (TCr) was nominally between −5 and +15 dB.

The onset of the target vocalization was drawn from an exponential distribution [72] (min: 1200 ms; mean: 4200 ms; max: 6100 ms). This strategy encouraged the monkey not to anticipate target onset [72]. On some trials, however, the vocalization chorus terminated without a target vocalization. These were *catch* trials.

The hearing-in-noise task was a detection task. A *hit* was when the target vocalization was presented and the monkey released the lever within 600 ms of target onset (i.e., the response-time window). A *miss* was when the vocalization was presented but the monkey did not release the lever within the response-time window. A *false alarm* occurred when the monkey released the lever when the target vocalization was not presented. A *correct rejection* was when the monkey held onto the lever throughout a catch trial. Monkeys were rewarded on hit and correct-rejection trials.

### 2.6. Data analysis

To determine whether vlPFC activity contained information that correlated with different TCr values (−5, 0, 5, 10, and 15 dB) and the monkey’s choices, we conducted two classification analyses on the µECoG signal. In one classifier, we test the degree to which we could decode different TCr values. In this analysis, we used µECoG signals generated from hit trials only of the hearing-in-noise task to ensure that we did not conflate behavioral choice with TCr. In the second classifier, we tested whether a classifier could discriminate between distributions of µECoG signals that were separated by the monkeys’ behavioral choices (hits, misses, false alarms, and, in a subset of sessions, correct rejections). We conducted these classification analyses on the µECoG broadband signal as well as the 4–8 Hz (theta band), 8–12 Hz (alpha band), 13–30 Hz (beta band), 30–90 Hz (gamma band), and 90–600 Hz (high frequency band) components of this signal.

For each classification analysis [73-76], we constructed an N-dimensional population response vector that constituted the µECoG signals from a population of N different electrodes to R repetitions of S conditions (i.e., each TCr value or behavioral choice). We used a Support Vector Machine to classify these data, and each classification underwent a k-fold cross-validation procedure. This procedure divided the training set of µECoG data into k smaller subset (i.e., folds) and, in an iterative fashion, one subset was tested by a model trained on the remaining k-1 subsets.

Because different numbers of trials might have occurred for different conditions (e.g., each TCr value), we subsampled our data to ensure that we had the same number of trials for each condition [77]. For each electrode, we calculated the mean µECoG amplitude and its variance that was generated over the entire duration of the target vocalization or the equivalent period prior to lever release for false alarm trials. To control for potential bias due to electrode-by-electrode differences in the µECoG signals, we z-scored the µECoG signal, relative to a 750-ms baseline period preceding chorus onset.

To test how well the µECoG signals discriminated between the different TCr values or the monkey’s choices, we implemented a linear-readout procedure. Because both the TCr and the choice classifications had more than two conditions, we implemented a “one-versus-all” classification. In this method, we built a classifier for each TCr value and trained each of them to discriminate between one particular TCr value versus all of the remaining TCr values. Using the test data, we identified the classifier that maximized performance and report average performance across TCr values. We report the fraction of times that the test data was classified correctly and report average performance over 500 different instantiations of this one-versus-all classification. An analogous procedure was conducted to analyze choice. All classifiers were constructed using a Support Vector Machine procedure that was implemented in the MATLAB programming environment and used the LibSVM library [78] with a coarse-grained search that optimized the LibSVM parameters (e.g., cost function) for each classifier.

Because single-electrode quality varied on a day-by-day basis (see Figure 7), we instituted an algorithm to identify the individual electrodes that would be used in the classifier analysis. For each electrode and for each test session, we conducted a classifier analysis that was analogous to that just described. However, in this analysis we just analyzed individual electrodes; i.e., N was always equal to 1. We then identified those electrodes in the top 20^th^ percentile of classification accuracy and used these to construct the N ≥1 dimensional population response vector.

Chance performance for each classifier was measured by a bootstrap procedure in which we first randomized the relationship between the data and its label (i.e., the TCr value or the particular choice) and then conducted a classification analysis analogous to that described above. Estimates of chance performances were calculated by independently repeating this process 100 times. Decoding performance was considered to be different than chance when the 95% confidence intervals of the actual decoding accuracy did not overlap with the confidence intervals of this bootstrapped distribution.

**Figure 7.**
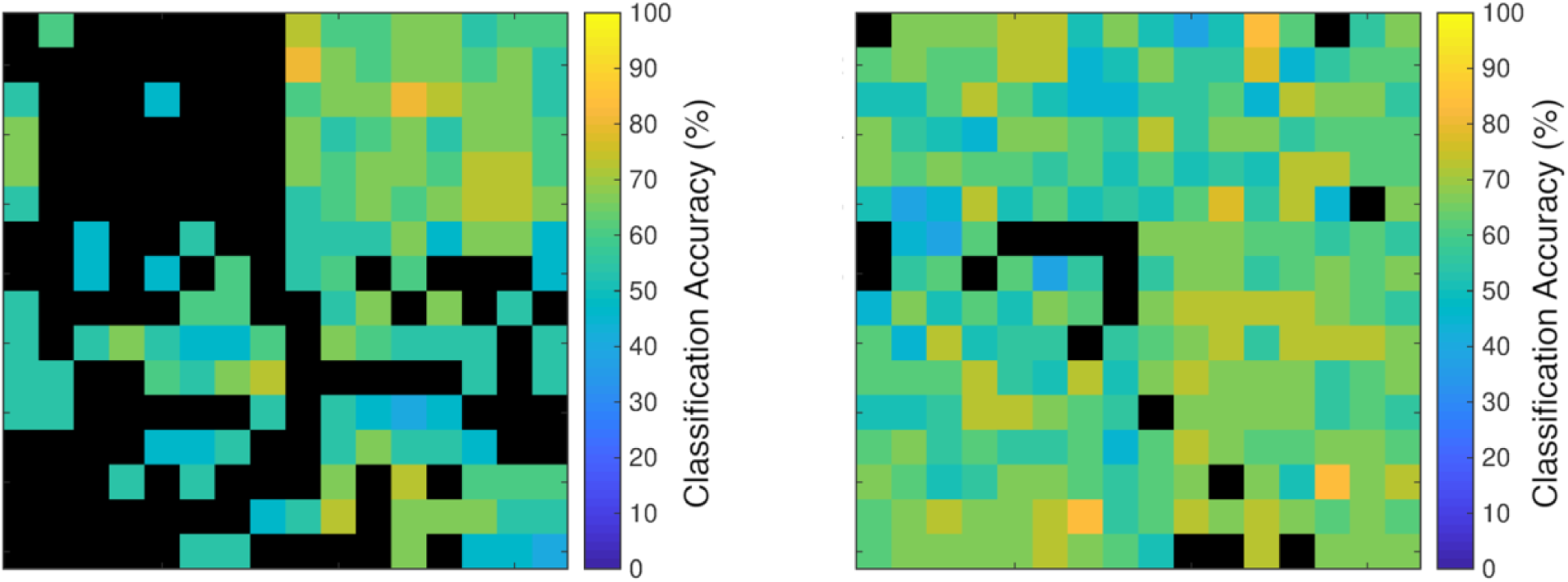
Individual channel decoding accuracy. Single-channel decoding (classification) accuracy of the μECoG array (left) 5 months and (right) 11 months after implantation. Each square represents the classification decoding accuracy of a single electrode; see scale legend next to each panel. In this figure, we show the accuracy of a classifier trained to decode three different behavioral choices.

## 3. Results

The µECoG electrode array was implanted in June 2017. The recordings for this study began in November 2017 and continued for ∼6 months. As of early 2019, the electrode was still implanted and was still viable.

As noted in Materials and Methods, there was tremendous day-to-day variability in the decoding of individual electrodes. This variability can be seen in Figure 7, which depicts the single-channel decoding capacity of each µECoG electrode after 5 months of implantation and after 11 months of implantation. As can be seen, this variability was not stationary: the decoding pattern from 5-months post-implantation was quite different than that seen at 11 months. We could not identify a trivial cause for this variability, such as fluctuations in the impedances of the electrodes. Because of this heterogeneity, if we “blindly” decoded the entire array, it did not perform better than chance. Instead, after calculating the single-electrode decoding capacity (Figure 7), we sorted the channels by their decoding capacity and selected only those electrodes in the top 20^th^ percentile for further analysis.

In our first analysis, we decoded TCr as a function of the number of electrode channels and as a function of µECoG frequency band. Depending on the particular session, the decoding of N=1 channels was either at chance levels (i.e., 20% due to the 5 TCr values [-5, 0, 5, 10, and 15 dB]) or above (Figure 8). However, as N>1, decoding accuracy quickly improved and was significantly above chance. When we decoded N=16 channels simultaneously, decoding typically exceeded 70%. Nonetheless, although there was substantial variability across different recording sessions, decoding capacity was comparable across all tested frequency bands.

**Figure 8.**
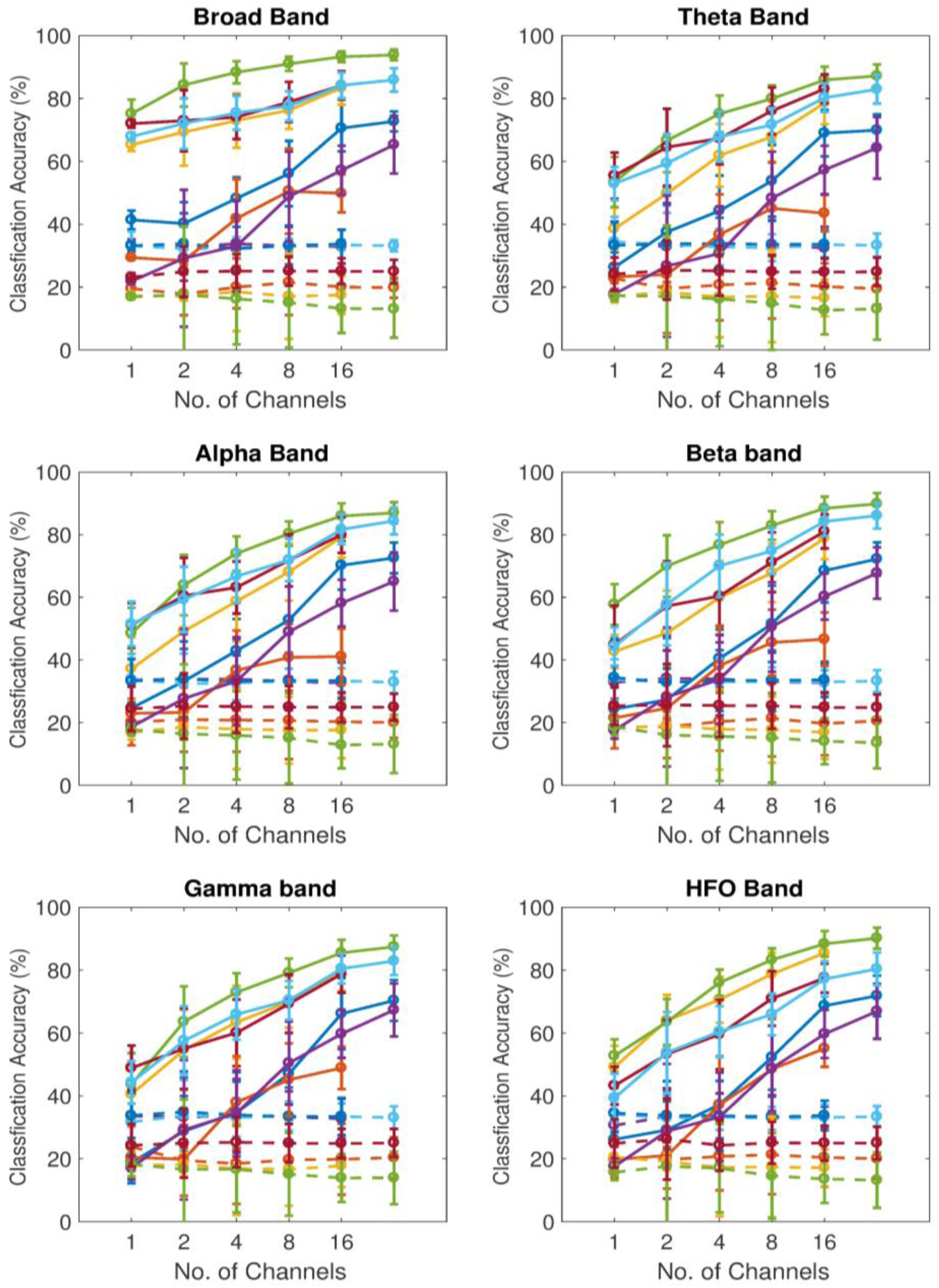
Population decoding accuracy of the µECoG array for TCr. Each panel shows the decoding (classification) accuracy of the μECoG array as a function of the number of simultaneously decoded channels. Solid lines show the mean decoding accuracy. Dotted lines represented mean classifier performance expected by chance (bootstrap randomization procedure). Error bars represent the standard deviation of the mean. Chance is theoretically at 20%. Different colors represent performance from different recording sessions. The different panels show decoding accuracy as a function of different neuronal frequency bands.

A somewhat different finding resulted from an analysis of behavioral choice as shown in Figure 9. Except for the broadband signal, when we decoded N=1 electrodes, the decoding accuracy ranged between chance (33% for 3 behavioral choices [hits versus misses versus false alarms] and 25% for 4 choices [hits versus misses versus false alarms versus correct rejections]) and ∼60%. Interestingly, for the broadband signal, decoding accuracy was ∼70% for N=1 electrodes. With more electrodes, decoding accuracy increased significantly and averaged ∼90% at N=16 electrodes. Although the single-channel decode of the broadband signal exceeded the other frequency bands, this advantage was minimized when we simultaneously decoded N=16 electrodes.

**Figure 9.**
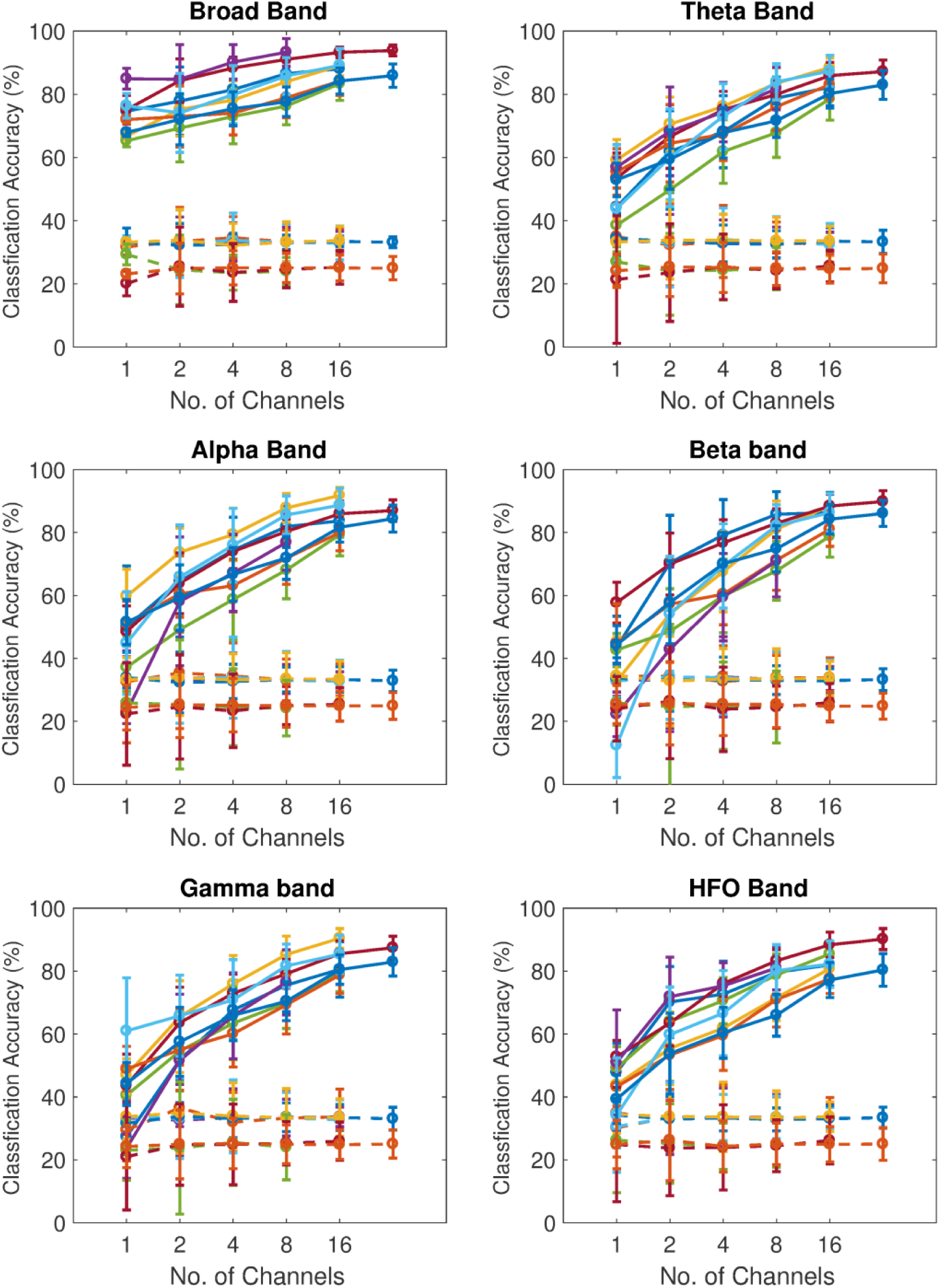
Population decoding accuracy of the µECoG array for behavioral outcome. In this data set, in different sessions, we decoded three or four different behavioral outcomes simultaneously; chance is 33.3% or 25%, respectively. The figure is formatted in the same manner as Figure 8.

Two inter-related differences are obvious when comparing this result with that of the TCr result in Figure 8. First, the day-by-day variability in decoding accuracy was smaller for the behavioral decode than with the TCr decode. Second, the decode capacity at the largest number of electrodes (N=16) was consistently better for the behavioral decode than for the TCr decode.

This session-by-session variability for both the TCr and behavioral decoding raises the obvious question of whether this reflects degradation of the µECoG electrode array over time or whether it represents some other idiosyncratic fluctuations in the electrode array. To address this question, we plot the decoding capacity of the array for behavioral choice as a function of time from implantation (Figure 10) for the broadband and gamma-band data; some of this data is a reformatting of the data shown in Figure 9. Over an approximate 1-year period (relative to surgical implantation), we did not identify any monotonic temporal degradation of decoding accuracy.

**Figure 10.**
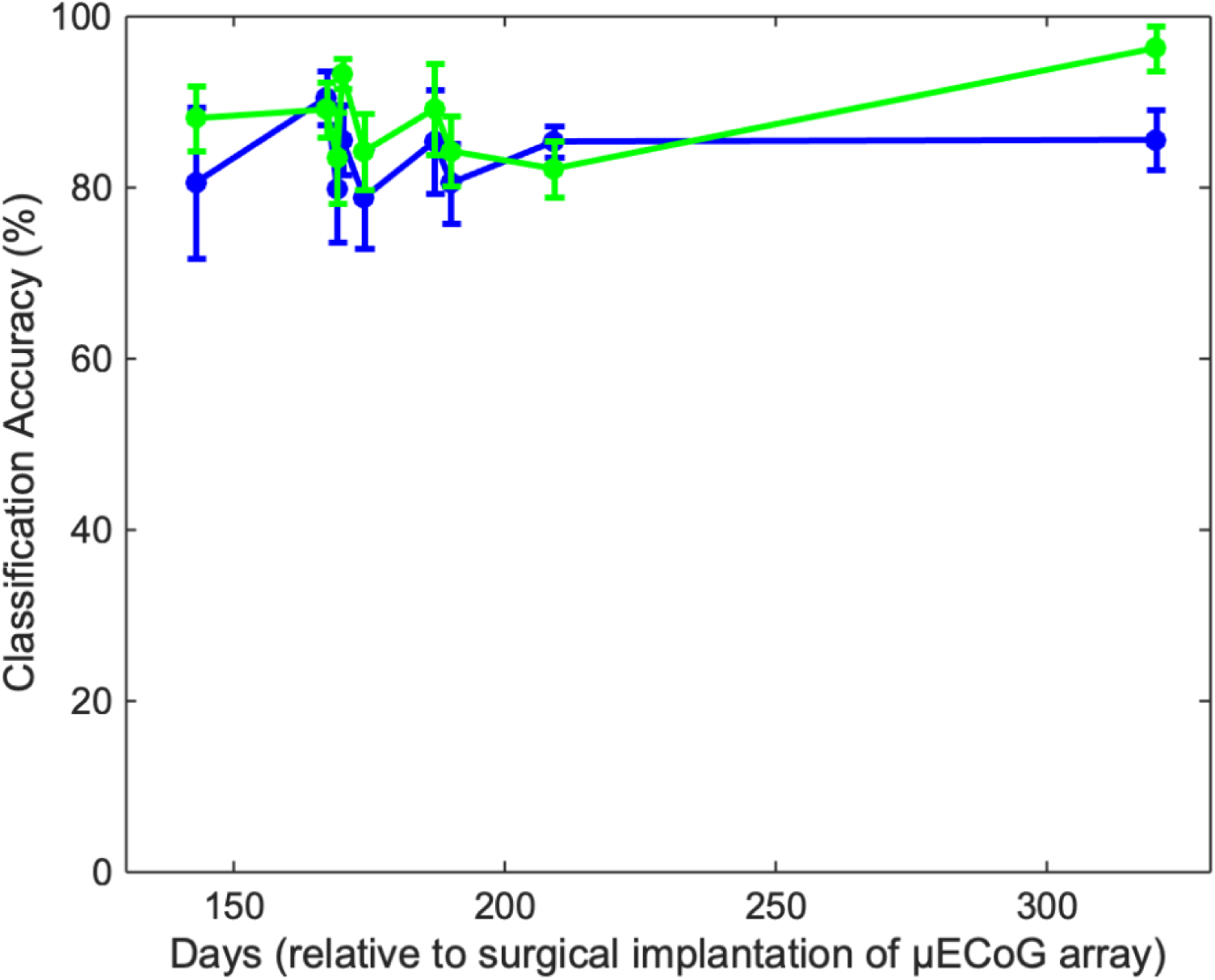
Population decoding accuracy the μECoG array as a function of time. The broadband (green) and gamma-band (blue) choice data from Figure 9. Classification accuracy was from 16 simultaneously decoded channels. Error bars represent the standard deviation of the mean.

In our final analysis, we asked whether the spatial organization of the electrode arrays affected decoding accuracy. This question was motivated by the observation that with as few as N=2 randomly selected electrodes, we could decode TCr and behavior at better than chance levels. Given this observation, is there any utility in developing a denser µECoG array with hundreds of electrodes?

To explore this question, we conducted another series of analyses in which we sampled over different spatial extents of the array. We first identified all of the electrodes that had classification accuracy in the top 20^th^ percentile. Next, we picked N=4 electrodes so that these channels were evenly distributed across the entire electrode array (i.e., “low density”) and tested the decoding accuracy of this low-density electrode subset. Our “mid density” array was an N=16 subset of electrodes that evenly tiled the electrode array. Finally, we repeated this process for all channels that were in the 20^th^ percentile of classification accuracy (“high density”). We found that decoding accuracy was a function of electrode density (Figure 11). As the sampling density increased, decoding accuracy improved and, in some cases, improved by almost 25%. This improvement was seen across all frequency bands.

**Figure 11.**
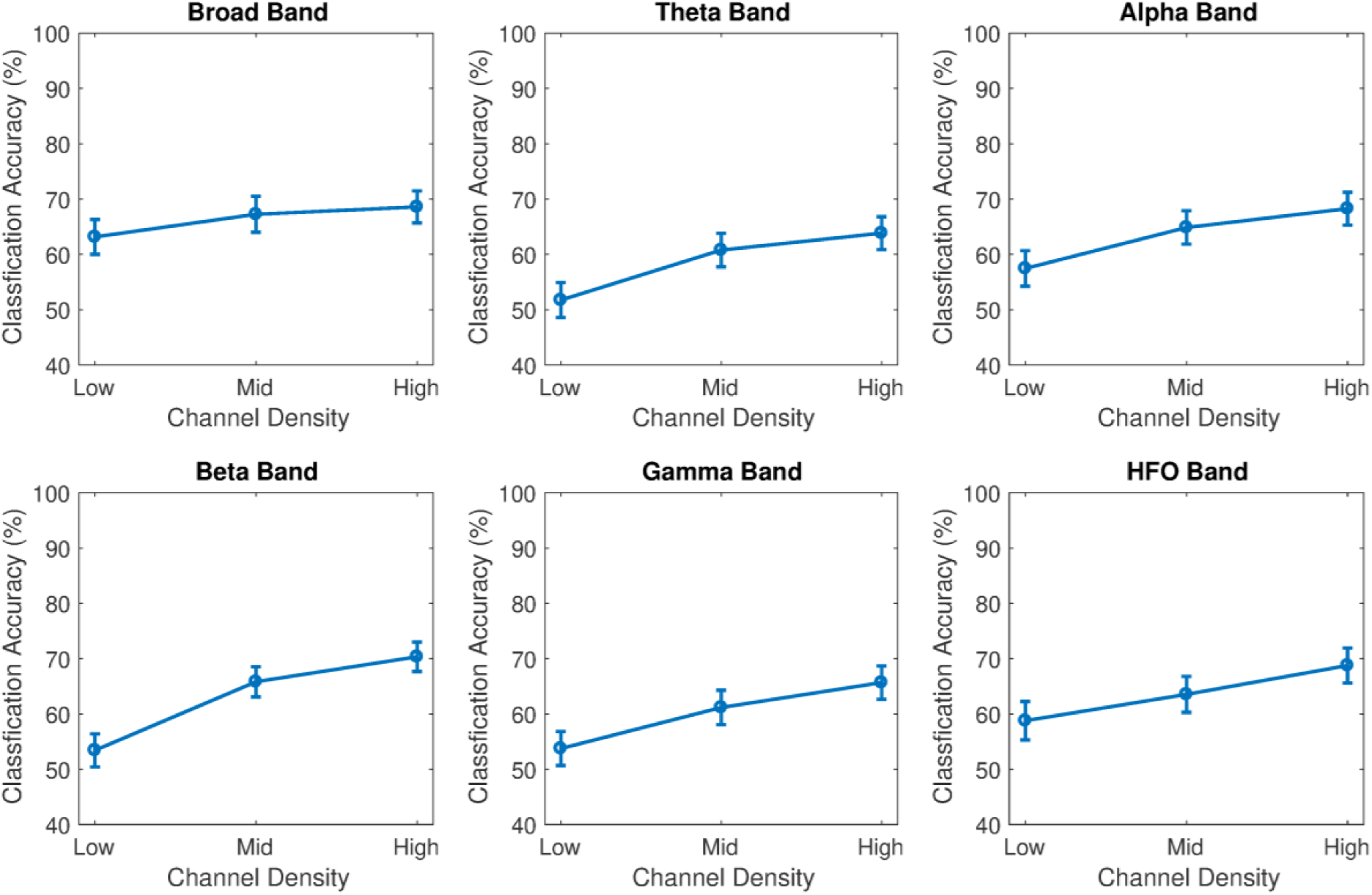
Decoding (classification) accuracy as a function of electrode density. Each panel shows the mean decoding accuracy for behavioral choices of the μECoG array as a function of channel density; error bars represent the standard deviation of the mean. Low-density classification corresponded only to four electrodes that had classification accuracy in the top 20^th^ percentile, whereas high density corresponded to all electrodes that had classification accuracy in the top 20^th^ percentile. The different panels show decoding accuracy as a function of different neuronal frequency bands.

## 4. Discussion

We developed a modular recording system that can scale to record larger channel counts without increasing the footprint of the implant. Utilizing silicone molding, we combined the sensing area of three thin-film sub-arrays to form a uniform high-resolution µECoG array. This array contains 294 channels and covered 10.4 × 11 mm^2^ without compromising flexibility of the device or the miniature footprint of the chamber. The modular approach not only allows customizations of electrode size, shape, and density for the targeted brain area but also allows the acquisition interface to expand vertically instead of horizontally, which minimizes the implant footprint. In addition, this modular approach could expand and scale to record over a thousand channels with larger brain coverage by stacking more PCBs and molding more electrodes together.

We also developed a design approach for custom 3D-printed titanium recording chambers that are watertight, optimized for surface arrays, and are individually mated to the skull of each animal and recording target. Unlike penetrating microdrives that rely on penetrating through a combination of silicone grease, silicone sealant, and Silastic membrane to prevent fluid ingress [79], surface electrodes typically have a flexible cable, which is thin but wide, which cannot penetrate through these sealant layers. To achieve a watertight seal, we developed a molding technique that generates a highly customizable silicone rubber wall that compresses and deforms around the electrode cable to seal the electrode entry point. We also created a silicone gasket to seal the gaps between the chamber cap and base. This design approach can be easily applied to all surface array implants and allows the PCBs within the chamber to be upgradable to add additional capabilities such as recording and stimulation.

When we used the µECoG array to decode different task parameters (Figures 7-10), we had two major findings. First, there was substantial day-to-day variability in the decoding capacity of each electrode. Nonetheless, when we examined multiple electrodes simultaneously, we found that we achieved significant and almost near-perfect decoding accuracy. Second, independent of the channel count, we found that a high-density array of µECoG electrodes outperformed lower density configurations. This finding supports future work aimed at developing chronic high-density arrays.

From a neuroscience perspective, this study contributes substantially to our knowledge of the vlPFC. First, it is not clear what information is encoded in the spiking activity of vlPFC neurons. Is it low-level stimulus information or higher-level perceptual information [60, 80-83]? Here, we found that the µECoG signals, which are surface electrical brain potentials, coded both types of information: sensory (TCr) and perceptual (choice). This rich and diverse quality of information has tremendous potential for future studies that would utilize vlPFC, or more generally PFC, µECoG signals, as part of a brain-machine interface.

Finally, even though µECoG signals reflect, to some degree, pooled electrical activity, the individual channels did not accurately decode TCr or choice. We only had near-perfect decoding accuracy when we simultaneously decoded multiple channels in a high-density regime across the entire array; a phenomenon that we found across multiple frequency bands. This suggests that vlPFC contains a rich and distributed neuronal code, which exists across multiple temporal scales, for task-related behavior.

## 5. Conclusion

We developed a new high-resolution surface electrode array that can be scaled to cover larger cortical areas without increasing the chamber footprint. Further, we demonstrated the utility and robustness of this high-resolution chronic µECoG array by finding that both sensory and perceptual information can be decoded from vlPFC µECoG signals. Independent of the number of electrodes used for decoding, we found that a high-density array of µECoG electrodes outperformed lower density configurations. This finding supports future work aimed at developing chronic high-density arrays.

## 6. Acknowledgements

This work was supported by grant W911NF-14-1-0173 from the Army Research Office (ARO) and grants from the NIDCD.

---

**Supplement Fig. 1.**
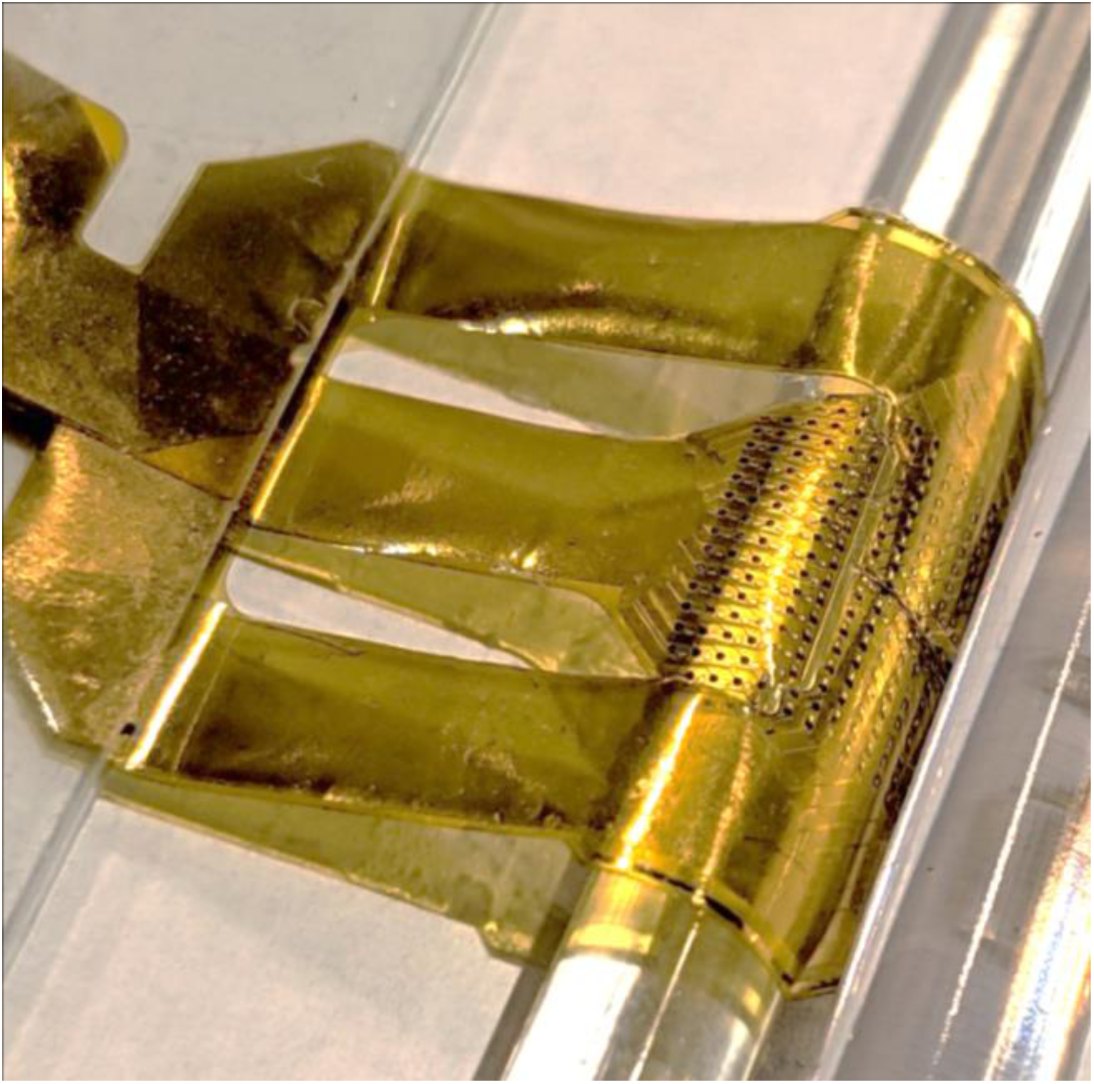
Electrode array bending test. The molded electrode array was not damaged and remained functional after hundreds of bend cycles on a glass rod. The radius of the glass rod was 2.5mm.

**Supplement Fig. 2.**
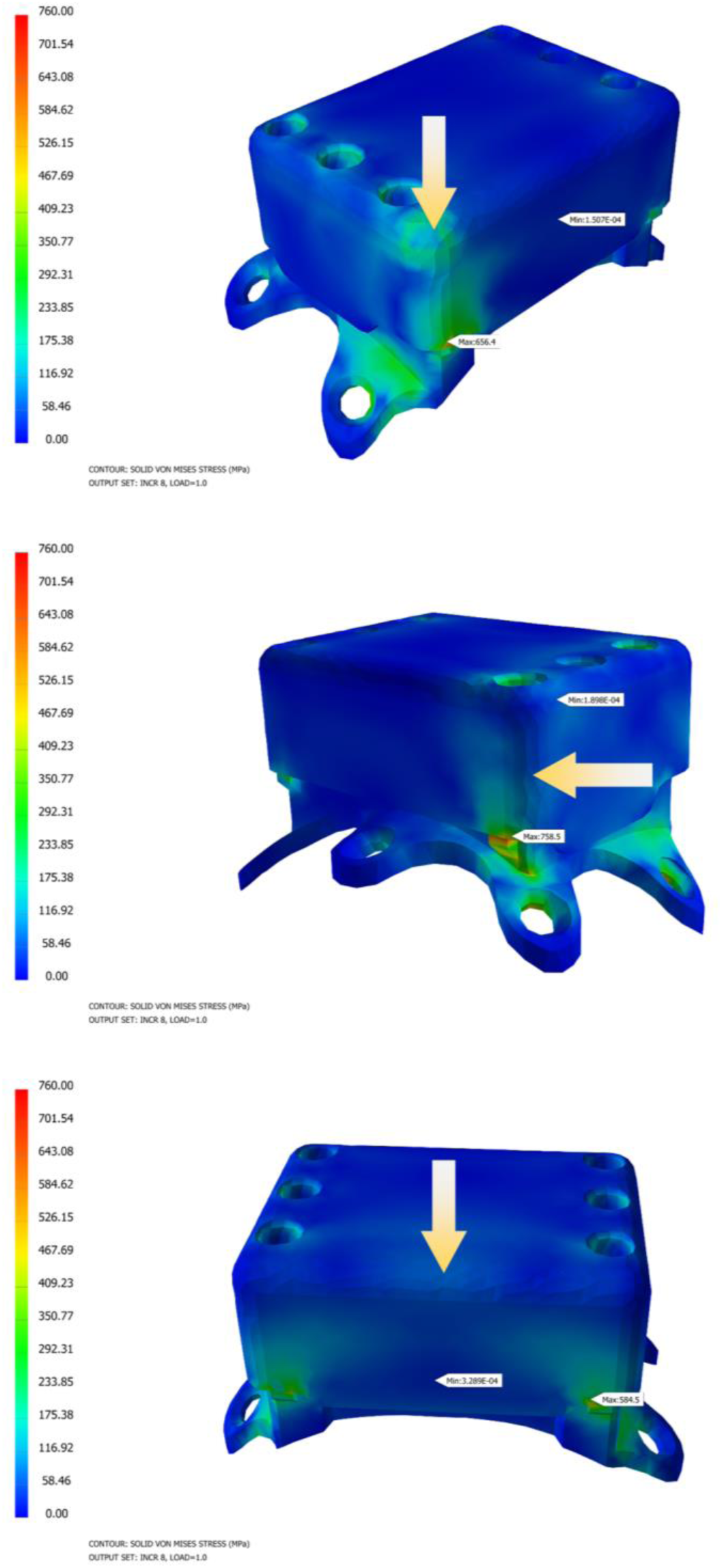
Mechanical simulations of chamber impact force from impacts at three alternate angles. Yellow arrows point to the entire edge or corner where the impact force of ∼4300N was applied. The maximum pressure remains below the yield strength limit of Ti.

**Supplement Fig. 3.**
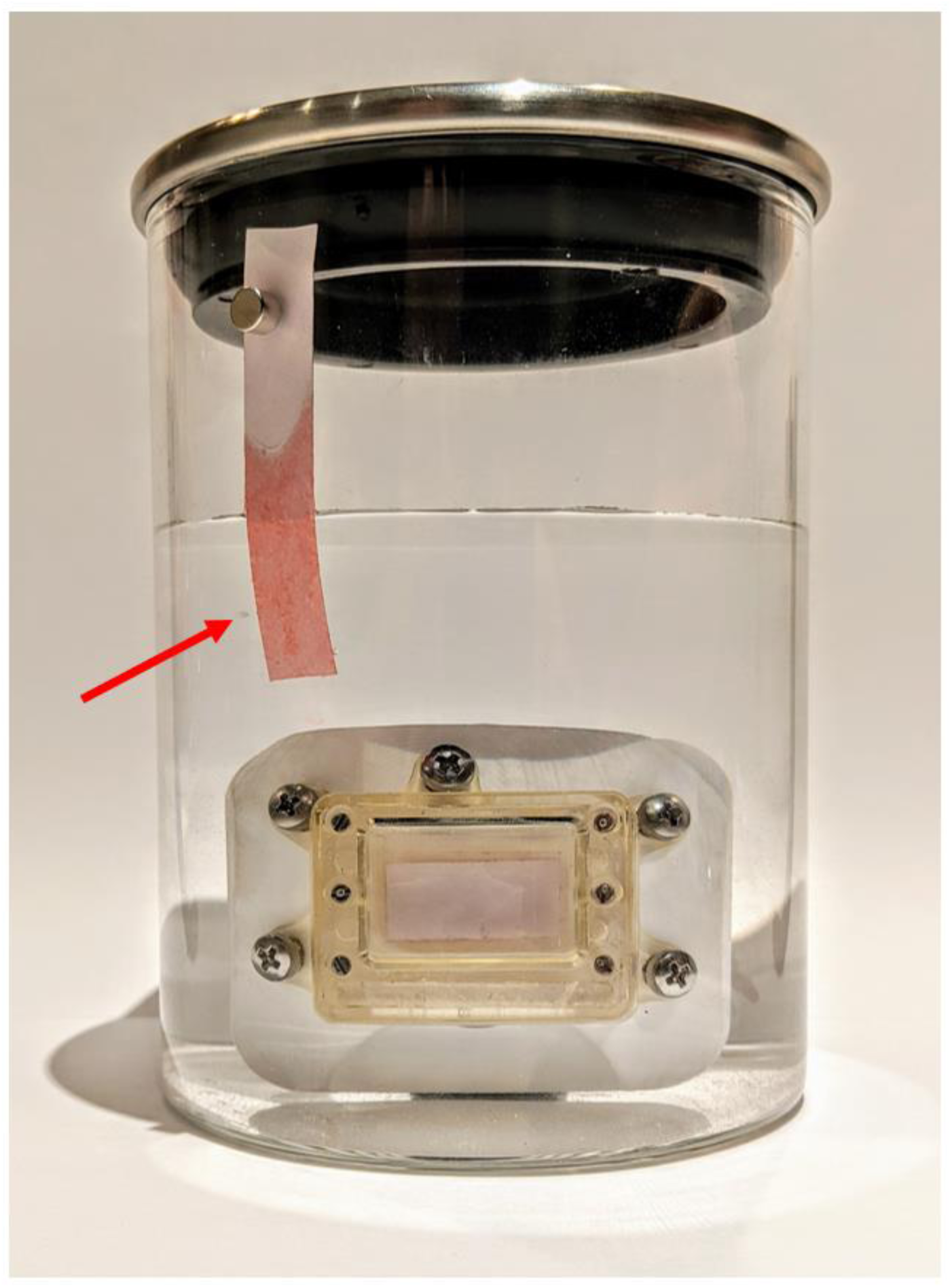
Recording chamber water resistance testing. A 3D-printed clear chamber was assembled with a water-contact indicator strip (3M 5559) and soaked in PBS. The strip showed no-sign of water contact over two weeks of soaking (i.e. stayed white/pink). A comparison strip outside the soaking jar (highlighted by the red arrow) shows the color (red) the strip changes to when the tape is exposed to liquid water.

**Supplement Fig. 4.**
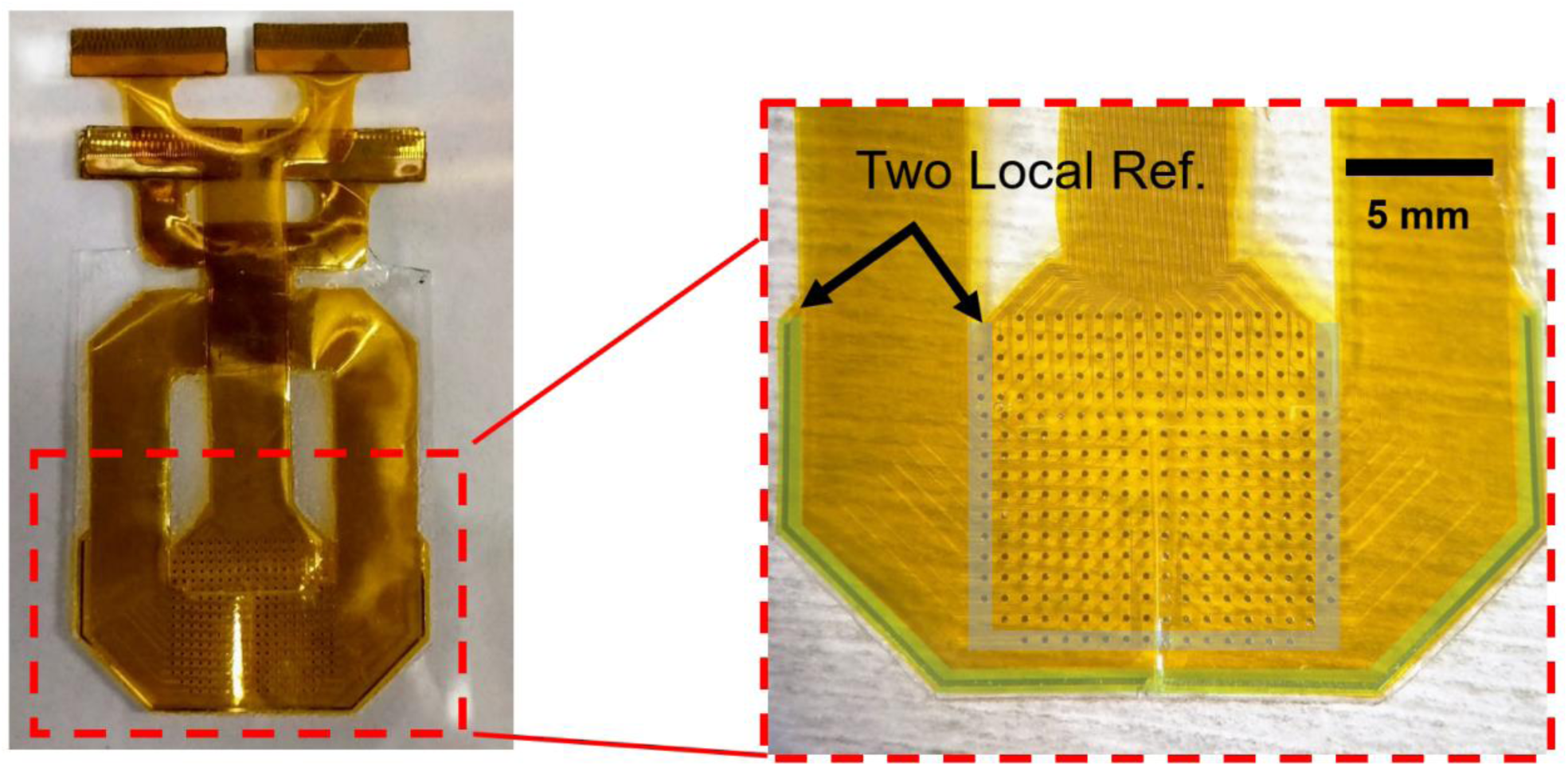
Electrode array local reference. The electrode array included two local references. First, an outer, large-area contact (highlighted in green), which was used as the reference in all recordings. Second, 40 electrode contacts that could not be recorded by the Open Ephys system due to space and channel limitations (highlighted in blue), were shorted together in the PCBs and recorded as a single reference channel. This channel can be used to optionally re-reference the recorded signals.

